# Differentiation of vegetative cells into spores: a kinetic model applied to *Bacillus subtilis*

**DOI:** 10.1101/309617

**Authors:** Emilie Gauvry, Anne-Gabrielle Mathot, Olivier Couvert, Ivan Leguérinel, Matthieu Jules, Louis Coroller

**Affiliations:** Université de Brest, EA 3882, Laboratoire Universitaire de Biodiversité et Ecologie Microbienne, UMT Spore-Risk, IUT, 2 rue de l’université, 29334 Quimper; Université de Brest, EA 3882, Laboratoire Universitaire de Biodiversité et Ecologie Microbienne, IBSAM, UMT Spore-Risk, 6 rue de l’université, 29334 Quimper; Université de Brest, EA 3882, Laboratoire Universitaire de Biodiversité et Ecologie Microbienne, IBSAM, UMT Spore-Risk, ESIAB, 2 rue de l’université, 29334 Quimper.; icalis Institute, INRA, AgroParisTech, Université Paris-Saclay, 78350 Jouy-en-Josas, France

## Abstract

Bacterial spores are formed within vegetative cells as thick-walled bodies resistant to physical and chemical treatments which allow the persistence and dissemination of the bacterial species. Spore-forming bacteria are natural contaminants of food raw materials and sporulation can occur in many environments from farm to fork. In order to predict spore formation over time, we developed a model that describes both the kinetics of growth and the differentiation of vegetative cells into spores. The model includes a classical growth model with the addition of only two sporulation-specific parameters: the probability of each vegetative cell to sporulate, and the time needed to form a spore once the cell is committed to sporulation. The growth-sporulation model was evaluated using the spore-forming, Gram positive bacterium, *Bacillus subtilis* and the biological meaning of the sporulation-specific parameters was validated using a derivative strain that produces the green fluorescent protein as a marker of sporulation initiation. The model accurately describes the growth and the sporulation kinetics in different environmental conditions and further provides valuable, physiological information on the temporal abilities of vegetative cells to differentiate into spores.

**Importance:** The growth-sporulation model we developed accurately describes growth and sporulation kinetics. It describes the progressive transition from vegetative cells to spores with sporulation parameters which are meaningful and relevant to the sporulation process. The first parameter is the mean time required for a vegetative cell to differentiate into a spore (i.e. the duration of the sporulation process). The second sporulation parameter is the probability of each vegetative cell forming a spore over time. This parameter assesses how efficient the sporulation process is, how fast vegetative cells sporulate and how synchronous the bacterial population is for sporulation. The model constitutes a very interesting tool to describe the growth and the sporulation kinetics in different environmental conditions and it provides qualitative information on the sporulation of a bacterial population over time.

## Introduction

Spore-forming bacteria are common contaminants of food, and represent the major source of food poisoning and food spoilage (1, 2). The aim for industrials is to prevent contamination of foods by bacterial cells under their vegetative or sporulated forms. To do so, it is necessary to target and control the different steps of the life cycle of these microorganisms. Bacterial cells under their vegetative or sporulated forms can be found in the environment and thereby can be natural contaminant of raw materials. The spore-formers display many physiological and enzymatic capacities. The spores are commonly resistant to physical and chemical treatments applied in the food industry. On the contrary, vegetative cells are sensitive but they can grow, produce degradative enzymes or toxins, form biofilms and differentiate into resistant spores as observed in milk powder processes (3–5).

In order to control the occurrence of spore-formers in foods and in the food industry, it is necessary to prevent the growth and the sporulation of these microorganisms. A better understanding of the ecological niches of spore-formers can help preventing raw material contamination (6, 7). The sporulation leads to an increase of the spores yield in foods and the sporulation conditions affect the quantity and the resistance properties of spores to subsequent chemical or thermal treatments (8, 9). The tools of predictive microbiology can help preventing the different bacterial processes thanks to mathematical models. The bacterial growth can be predicted over time and according to environmental factors (10–12). And some models exist to predict the resistance of spore according to chemical and physical treatments also (13–16). However, the sporulation process has been largely ignored in predictive microbiology.

Mechanistic, knowledge-based models of sporulation have been proposed to describe the decision-making process of sporulation initiation at the cellular and molecular levels in response to environmental stimuli (17–19). These models are complex because they require numerous parameters, which for most of them cannot be experimentally evaluated in industrially relevant conditions. Alternatively, empirical, phenomenological models of sporulation were proposed to describe the evolution of spore counts over time, as they are simpler to use than mechanistic models. However, empirical models do not take into account the fact that sporulation is a differentiation process of vegetative cells into spores (20, 21), while growth and sporulation are well-known to be interdependent physiological processes (22).

Sporulation occurs following different signals such as nutrient starvation and communication molecules of quorum sensing, that require previous bacterial growth. After signal sensing (23) the sporulation starts with the activation by phosphorylation of the master regulator Spo0A until a given threshold of Spo0A~P. Once this threshold is reached, the activated master regulator activates the early sporulation genes such as *spoIIAA* in the pre-divisional cell and triggers the asymmetric division to form the mother-cell and the forespore (24). The sporulation process continues according to a sequential process involving different transcription factors specific to the mother cell (σ^E^ and *σ*^K^) and to the forespore (σ^F^ and *σ*^G^) until the formation of a mature spore.

The objectives of this work were to develop a model that (i) describes the sporulation kinetics from the growth kinetics of vegetative cells and (ii) can be used to predict sporulation in industrially relevant conditions. The identification of the model parameters required to assess the temporal heterogeneity of the sporulation of the vegetative population over time, the time that the vegetative cells needed to complete the sporulation process and the sporulation efficiency. To assess the biological meaning of the sporulation parameters, the model of the Gram positive bacteria, *Bacillus subtilis* was used in combination with a fluorescent reporter of sporulation initiation (P *_spoIIAA_*-*gfp*).

## Results

### Model development and experimental strategy

A kinetic model associating the sporulation to the bacterial growth was developed. It describes the growth of vegetative cells with a classical logistic (Equation 1), and their differentiation into spores over time with two sporulation parameters: the probability of vegetative cells to sporulate over time and the time for each cell to form a mature spore *t_f_* (we assume that all vegetative cells need the same time to form a spore). The probability to sporulate was defined at the maximum population level by the proportion of vegetative cells (out of 20 cells in the exemple depicted in Figure 1a) which initiate the sporulation over time. At the cell level, this proportion accounts for the probability of each individual cell to sporulate over time (Figure 1b). This probability to sporulate evolves over time following a Gaussian distribution (Equation 3), which is described with three parameters (Figure 1b). The first parameter is the maximal probability to sporulate*P_max_* which accounts for the maximal proportion of vegetative cells that can sporulate in a given period of time. This parameter mainly influences the maximal concentration of spores to be produced. The second one is the time *t_max_* at which this maximal probability to sporulate is obtained, which has an impact on the time at which the first spores appear. The third parameter is the probability scattering which has an impact on the speed of appearance of spores over time.

**Figure 1.**
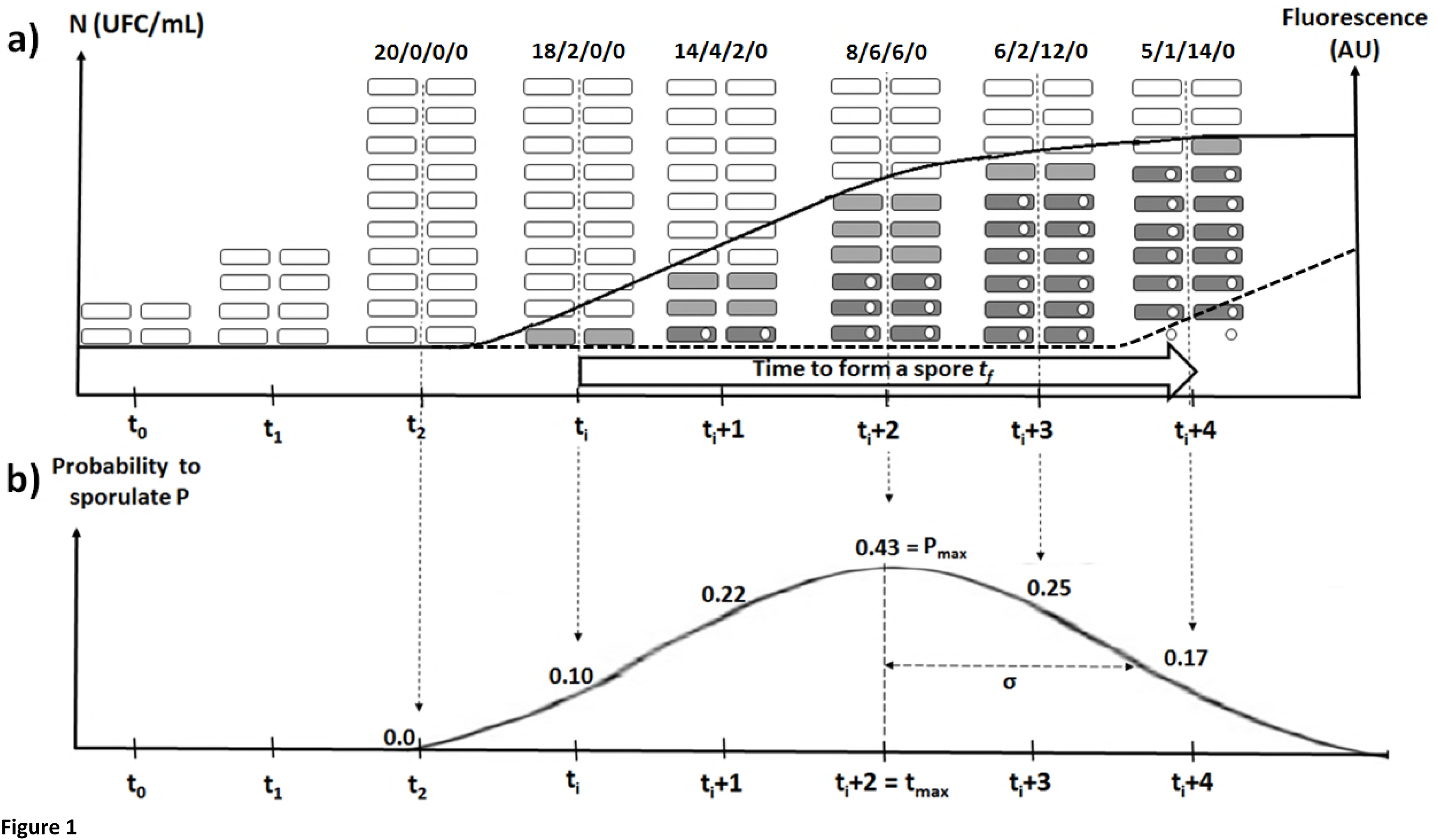
Schematic representation of the growth and sporulation model. The bacterial population of the strain P*_spoIIAA_ gfp* can be divided into four sub-populations (Figure 1a). Among the total cells (20 cells in this example), there are the vegetative cells not committed to sporulation (▭), the vegetative cells that initiate the sporulation process at each time of the culture 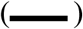 and produce GFP 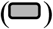, the vegetative cells already committed to sporulation (or sporulating cells) 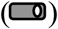 and the mature spores (o) defined as resistant cells in our study (○) with its corresponding curve 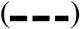. The proportion of each sub-population is given by the numbers separated by slashes. In this figure, the vegetative cells (4 cells are inoculated at time to) grow until they reach 20 cells at time t_2_ (following Equation 1). At each time of the culture, a given proportion of vegetative cells not committed to sporulation yet initiates the sporulation, what defines the probability to sporulate over time (Figure 1b). Once the sporulation is initiated, this process takes some time to achieve and form a mature spore, what defines the time to form a spore.

The experimental strategy developed to assess the sporulation parameters consisted in using a promoter fusion between the *gfp* gene and the promoter of the gene *spoIIA* (P_*spoIIA*_ *gfp*) as a reporter of the initiation of sporulation. We made the hypothesis that on average, each sporulating cell produces the same amount of GFP (*i.e.* they produce the same amount of fluorescence). Consequently, the increase of the fluorescence over time (right scale in Figure 1a) accounted for the increase of sporulating cells over time. The fluorescence and the concentration of sporulating cells evolved following a Gaussian distribution function (Equation 4). This allowed calculating the evolution of the probability to form as spore over time which evolves following the Gaussian density function (Equation 3 and Figure 1b). Ultimately, the time to form a spore was assessed (Equation 5) as the increase of fluorescence accounting for the increase of mature spores after the time to form a spore (dashed line in Figure 1a).

### Assessment of the growth and sporulation parameters of B. subtilis P_*spoIIAA*_ *gfp* at 27°C, 40°C and 49°C

The proposed models (Equations 1 and 4) accurately described the growth and sporulation kinetics. The qualities of fit for growth and sporulation models reached a global RMSE value of 0.90 ln (UFC/mL) for all conditions tested.

The growth and sporulation kinetics were not significantly different between the wild-type BSB1 and *P_spoIIAA_ gfp* strains for the three temperatures tested. The values of the likelihood ratio test were 8.37, 7.43 and 3.00 at 27 °C, 40°C and 49°C respectively, *i.e.* inferior to 15.51 (α< 5%). This allowed the wild-type strain to be used as a background to compute the fluorescence related to the production of GFP by strain P_*spoIIAA*_ *gfp*.

At 40 °C, the lag time was of 1.6 h, the growth rate was 1.61 h^-1^ (Figure 2f and Table 1) and cells reached a maximal concentration of 3.8 × 10^8^ CFU/mL at 10 hours of culture. The fluorescence of strain P*_spoIIAA_ gfp* increased with growth until it reached a maximal value *F_max_* of 5.13 × 10^4^ AU at 50 hours of culture. The maximal accumulation of fluorescence per unit of time was obtained at 36.7 h of incubation (*t_max_*) and with a standard deviation of 10.4 h (Figure 2d and 2e, and Table 1). The sporulation kinetics displayed a first phase of abrupt appearance of almost 10^3^ CFU/mL and a second phase with a more gradual appearance of spores over time. These two phases were correctly described by the predicted kinetics. The maximal concentration of spores was 4.86 × 10^5^ CFU/mL (Figure 2f) and was directly linked to the maximal sporulation probability P*_max_* which was estimated at 2.4 × 10^-2^ (Table 1). The use of the model allowed computing a time to see the first spore at 9.0 h of culture which was consistent with experimental observations. Indeed, the time needed to obtain the first 10 spores per milliliter (corresponding to the detection limit) was at 12 h of culture. Lastly, the time to form a spore was estimated at 7.0 h of culture which was consistent with previous findings (25).

**Figure 2.**
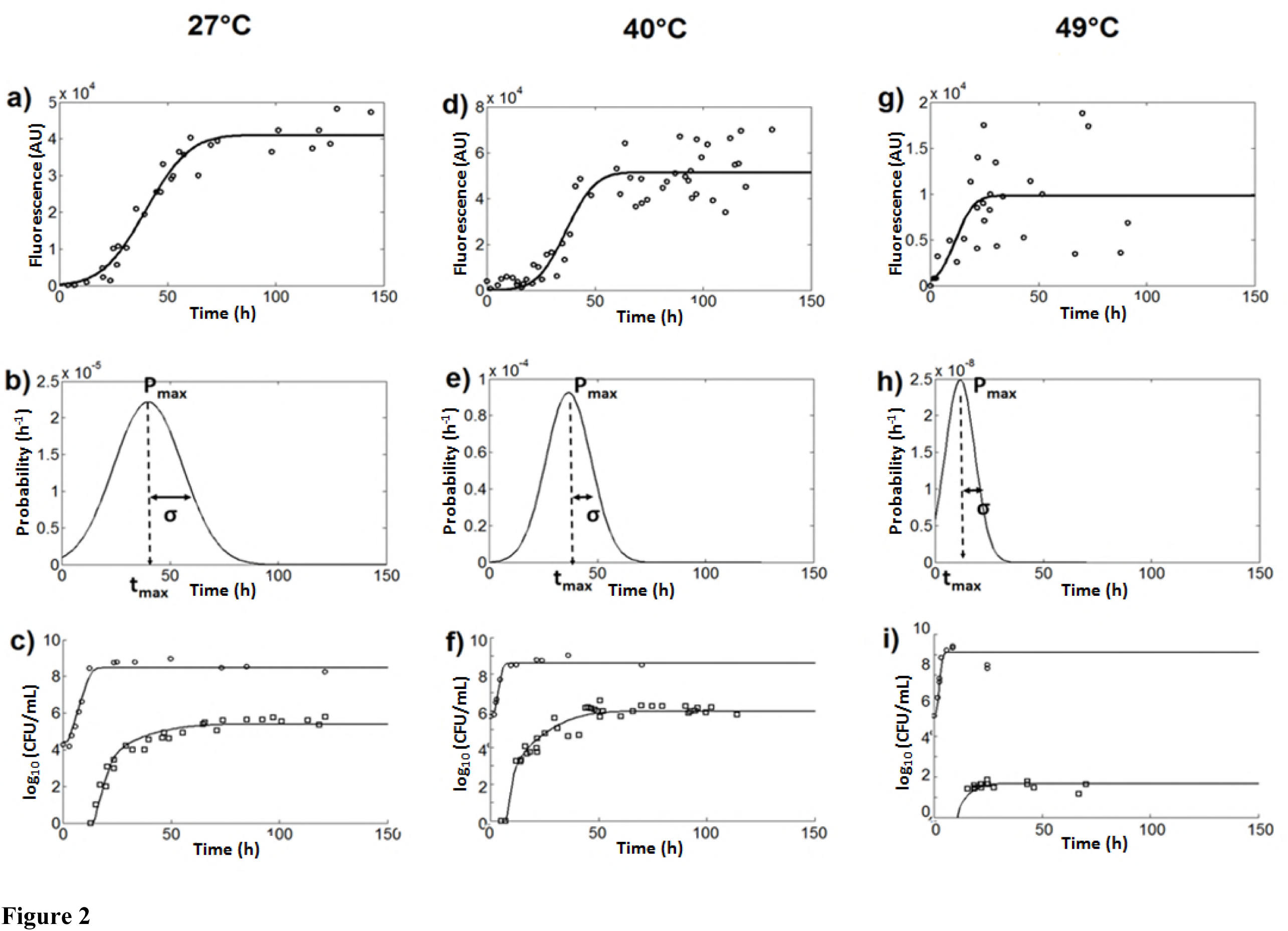
Fluorescence, growth and sporulation kinetics of *B. subtilis* at 27°C (a, b and c), 40°C (d, e and f) and 49°C (g, h and i). The values of fluorescence (o) were fitted with the normal density function (solid lines in a, d and g) and the corresponding probability densities (b, e and h) with the three sporulation parameters of Equation 6: P_max_, t_max_ and *σ*. The concentration of total cells (o) and the concentration of spores (⧠) over time were fitted with the growth sporulation model in equations 1 and 4 (in c, f and i).

**Table I.**
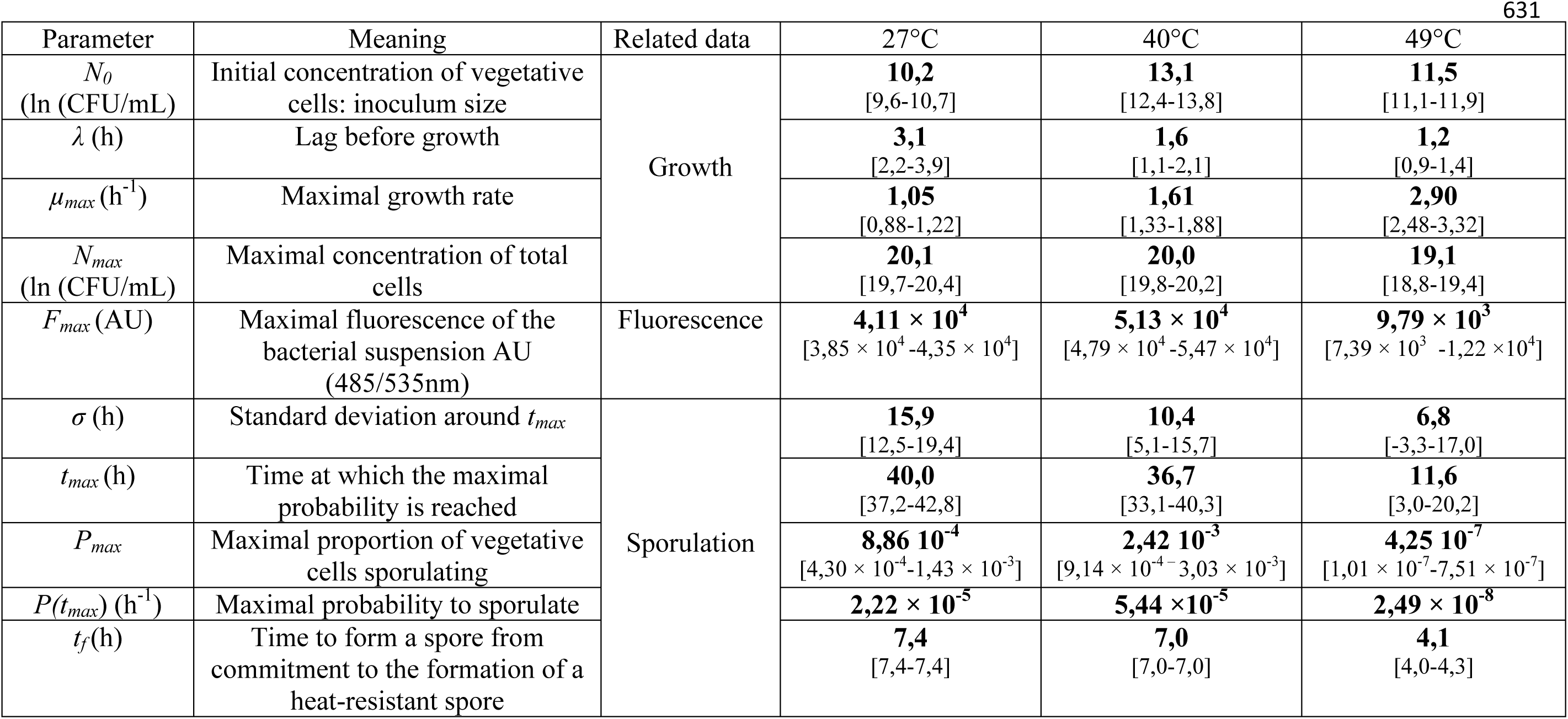
Estimations of the fluorescence, the growth and the sporulation parameters of *B. subtilis* at 27 °C, 40 °C and 49 °C. Values between brackets correspond to the confidence intervals (95%) of the estimates (bold values).

At 27 °C, the growth rate was reduced by 35% as compared to growth at 40°C, and the lag time was twice as high with *λ* values of 1.6 h and 3.1 h at 40 °C and 27 °C respectively (Table 1). The fluorescence evolved more gradually from 0 h to 70 h at 27 °C than at 40 °C (Figure 2a). This led to a more scattered probability of commitment to sporulation at 27 °C with a *σ* value of 15.9 h compared to 10.4 h at 40 °C (Figure 2b) which explains the gradual appearance of spores at 27 °C (Figure 2c). The maximal fluorescence was 20% lower at 27 °C than at 40°C, leading to the estimation that the maximal sporulation probability was about 3fold lower at 27 °C than at 40°C. Thus, this explains why the maximal concentration of spores was 4-fold lower at 27 °C compared to 40 °C. The time taken to form a spore was estimated at 7.4 hours at 27 °C (as for 40 °C).

At 49°C, the growth of *B. subtilis* was enhanced with a growth rate almost twice higher than at 40°C. However, the maximal concentrations of total cells and the lag time were not significantly different (Table 1). The GFP-related fluorescence was detected as soon as growth started, increased faster than at 40°C and the maximal fluorescence was 5 times lower than at 40°C (Figure 2g compared to Figure 2d). The concentration of spores was reduced by 20,000-fold at 49°C compared to 40°C but the maximal probability to commit to sporulation was only reduced by 2,000-fold. Thus, the maximal probability was not sufficient to explain the observed difference in the spore yield. The maximal probability was obtained 25.1 h sooner, when the concentration of cells was much lower at 49 °C than at 40 °C. Consequently, the maximal concentration of cells which were able to sporulate in the same time was also lower at 49 °C. Furthermore, the probability was less scattered with a standard deviation *σ* around *t_max_* of 6.8 h at 49 °C compared to 10.4 h at 40 °C (Figure 2h and e). The probability scattering had an impact on the temporal accumulation of sporulating cells. When the probability scattering was low, cells were able to sporulate in a shorter time frame which led to fewer cells that were able to sporulate over time. Lastly, the sporulation process was faster at 49 °C than at 40 °C with times required to form a heat-resistant spore (*t_f_*) which were estimated at 4.1 h and 7.0 h at 49 °C and 40 °C respectively.

## Discussion

### Theories and design of the model

The aim and the originality of this work were to develop a model that describes both the growth kinetics and the sporulation kinetics with parameters that account for the differentiation of vegetative cells into spores. The sporulation was precisely described using the two parameters related to the decision-making process of cells to sporulate and the time they need to complete the process.

The logistic model of growth (Equation 1) is largely used to describe the bacterial growth. It describes the growth kinetics with the lag before growth, the growth rate and the maximal concentration of total cells. Similarly, some models were developed to describe the sporulation kinetics with parameters such as the lag before the appearance of the first spores, the sporulation rate and the maximal concentration of spores. However these models dissociate the growth and the sporulation whereas these two bacterial processes are physiologically intertwined (26).This statement was supported by previous observations on other species of *Bacillus* as a correlation between the growth rate and the sporulation rate was found (20).

The decision-making process to sporulate was defined elsewhere at the cell level (27 – 29) and was translated at the population level by the probability to sporulate *P* in this study. The sporulation decision-making process of vegetative cells is directly linked to both the growth rate and the bacterial density (26) which evolve themselves over time following the growth kinetic. Thereby, we suggested that the probability to sporulate evolves over time also. This hypothesis is supported by recent works by (30) who showed that the time of sporulation (or the time at which the cells enter into sporulation) is heterogeneous among a bacterial population. For many biological processes, heterogeneity is the result of the multiscale organization of life as explained elsewhere (31). The heterogeneity of sporulation between cells can be explained at molecular and cellular levels by stochastic variations (32). The heterogeneity of sporulation over time can be explained because the sporulation depends on nutrient starvation which becomes increasingly severe over time, and depends on quorum sensing molecules that accumulate over time. Moreover, the sporulation heterogeneity also rises with the heterogeneity of other decision-making cell processes such as entry into competence, cannibalism or dormancy (33, 34) that delay the entry into sporulation. Ultimately, once the sporulation is initiated by vegetative cells, the process takes some hours to achieve until it forms a mature spore, which defines the second sporulation parameter *t_f_*.

### Quantitative and qualitative information are brought by the sporulation parameters

The growth-sporulation model allowed describing accurately the growth and sporulation kinetics and allowed computing the time to obtain the first spore in the culture, the speed of appearance of spores and the maximal concentration of spores. Altogether, this revealed that the sporulation was the most efficient at 40 °C as the first spores appeared sooner and the maximal concentration of spores was higher than at 49°C and 27°C. This model allowed describing various curves shapes of growth and sporulation kinetics (fast and low kinetics) and was even more accurate than previous sporulation models (20, 21) with lower RMSE values (Supplementary Table S1). In particular, these early models did not succeed in describing the smooth emergence of spores as observed at 40 °C and 27 °C. In some cases, the use of these early models led to aberrant estimations of the time needed to see the first spores and the maximal concentration of spores (Supplementary Figure S1). Moreover, this model is capable of describing the growth and sporulation kinetics of other microorganisms such as *B. subtilis* BSB1 and *Bacillus licheniformis* Ad 978 and in various environmental conditions (Supplementary Figures S1 and S2).

The sporulation parameters also bring information at the physiological level on the sporulation behavior of vegetative cells over time. The probability to sporulate over time is described with a Gaussian density function involving three parameters. The maximal probability P*_max_* to sporulate accounts for the sporulation efficiency and explains why the sporulation yield is much higher at 40°C and 27°C than at 49°C. The low proportions of cells which sporulated at 49 °C may be the result of the rapid physico-chemical degradation of the medium provoked by such a high temperature. A simple hypothesis is that the deterioration of the growth medium may alter the cell decision-making and consequently advantage or disadvantage certain physiological processes; this hypothesis is supported by the rapid cell decline observed at 49 °C (Figure 2i).

The probability scattering *σ* assesses how synchronous the bacterial population is for initiating sporulation. At 49°C, sporulation was synchrone whereas at 40°C sporulation was much more asynchrone, as observed by the sporulating population heterogeneity. . At least two hypotheses can explain this observation. First, the temperature affects the membrane fluidity by modifying its composition in fatty acids, which in turn is known to affect the activity of the sensors such as the histidine kinase KinA (35). Second, differentiation processes such as the entry into competence or the cannibalism are impacted by environmental factors. For instance, *B. subtilis* displays cannibalistic behavior at 40 °C but not at 45 °C (36). Consequently, we can reasonably assume that there are fewer differentiation opportunities at 49 °C than at 40°C, which leads to a lower sporulating population heterogeneity at 49°C.

Concomitantly with *σ*, the time *t_max_* at which *P_max_* is obtained allows assessing the time at which the first cell initiates the sporulation, which is mathematically obtained when the product of the probability to sporulate with the concentration of total cells (CFU/mL) is superior to 1 *i.e.* 1 sporulating cell per milliliter. Lastly, the time to form a spore *tf* brings information on the time needed to complete the sporulation process according to environmental conditions. As the growth and the sporulation share enzymatic machineries (37–39), the time to form a spore is likely to be correlated with the growth rate. This could explain why the sporulation completed faster at 49°C where bacterial cells grew faster than at 40°C and 27°C. Nevertheless, dedicated experiments are required to address this issue.

In summary, a kinetic model was developed to describe both growth and sporulation as a differentiation process from vegetative cells into spores. On the one hand, the model describes the growth with the classical logistic model of Kono modified by Rosso (40). On the other hand, the models can be used to describe the sporulation kinetics from the growth kinetics with parameters thatare specific to sporulation: the time to form a spore and the probability to form a spore over time. The biological meaning of the sporulation parameters was experimentally assessed, providing both quantitative and qualitative information at the physiological level on the sporulation process. The sporulation parameters revealed that at suboptimal sporulation temperatures (eg. 49°C), vegetative cells commit to sporulation more synchronously, in lower amounts and belatedly than at optimal temperature (eg. 40°C). In the literature, few data are available on the time needed to complete the sporulation process and on the temporal behavior of vegetative cells for sporulation, according to environmental conditions of culture. The procedure we set to experimentally estimate the sporulation parameters experimentally offers new opportunities to better assess and understand spore formation across environmental conditions.

## Materials and methods

### Biological material and strain storage

The prototrophic *B. subtilis* BSB1 strain, a *trp^+^* derivative of *B. subtilis* 168, was used in this work (41, 42). The BSB1 derivative strain carrying the *P_spoIIAA_gfp* transcriptional fusion was built by transformation of genomic DNA from strain AC699 (kindly provided by Arnaud Chastanet, Micalis Institute, Jouy-en-Josas, France) using natural competence. Strain AC699 is a RL2792 derivative of the PY79 *B. subtilis* strain (43) containing the *gfpmut2* gene under the control of the *spoIIAA* promoter (*amyE*∷P*_spoIIAA_ gfp* / cat), which is a marker of the early stage of sporulation and controls the initiation of sporulation. The transcription of this gene is not subject to intrinsic noise, which means that the heterogeneity of activation of this gene is not due to stochastic processes but is correlated to the sensing of the environment (44). The GFP_mut2_ is stable for 7 days and in a pH range of 5.0 to 10.0 (45 – 47).

Concerning the transformation procedure, *B. subtilis* was grown overnight on Luria Bertani plates, (Difco™, Becton, Dickinson and Company) at 37 °C. After incubation, a colony was re-suspended in MG1 medium composed of MG medium (2g/L (NH_4_)_2_SO_4_, 1 g/L Na_3_C_6_H_5_O_7_, 14 g/L K_2_HPO_4_ ,3H_2_O, 6 g/L KH_2_PO_4_, 0.5% Glucose and 15.6 mM MgSO_4_) with an added 0.025% casamino acids and 0.1%, yeast extract for 4 h 30 min at 37 °C under 200 rpm agitation. A 10-fold dilution was then carried out in MG2 composed of MG medium to which 0.012% casamino acids, 0.025% yeast extract, MgSO_4_ 25mM and Ca(NO_3_)_2_ 8mM had been added. The suspension was incubated for 1 h 30 min at 37 °C under 200 rpm agitation (48). 200 μL of the suspension in MG2 was added to 0.1 μL of genomic DNA extracted from strain AC699 with a High Pure PCR Template Extraction Kit (Roche Dignostics, Meylan, France) and incubated for 30 minutes at 37 °C. Clones were selected on LB containing 5 μg/mL of chloramphenicol after incubation for 24 h at 37 C. The inability of the P*_spoIIAA_ gfp* strain to degrade starch (as the reporter fusion is inserted in the *amyE* locus) was also verified on starch plates with iodine revelation.

Concerning the storage procedure of *B. subtilis* strains, each selected colony was isolated on LB plates and incubated overnight at 37 °C. A colony was re-suspended in Luria Bertani Broth, Miller (Difco™, Becton, Dickinson and Company) under 100 rpm agitation at 37 °C for 4 hours. From this pre-culture, a 100-fold dilution was performed in 100 mL of LB broth in flasks, in the same culture conditions for 3 hours. A second dilution was then performed in the same conditions. When the early stationary phase was reached after a 5-hour culture, glycerol was added to the bacterial suspension at a final concentration of 25 *%* w/w in cryovials. The bacterial cells in cryovials were stored at -80 °C.

### Monitoring the kinetics of growth, sporulation and fluorescence

Vegetative cells were inoculated from the cryovials at an initial concentration of 1000 CFU/mL in 250 mL flasks filled with 100 mL LB broth, supplemented with sporulation salts (49). Bacterial cultures were performed under 100 rpm agitation, at 40 °C, which is close to the optimal growth temperature, and at two suboptimal temperatures for growth and sporulation (27 °C and 49 °C). The incubation was performed in darkness to prevent excitation and degradation of the GFP produced by the strain P*_spoIIAA_ gfp*.

The growth kinetics were monitored by pouring 1 mL of the relevant dilution into nutrient agar (Biokar Diagnostics, Beauvais, France). Enumeration of colonies was performed after incubation of the plates for 24 hours at 37 °C (ISO 7218). Sporulation was monitored by enumerating cells resistant to a 10-minute heat treatment at 80 °C. The heat treatment was applied to the suspension samples using the capillary method (8).

The green fluorescence emitted by the total suspensions of the wild-type BSB1 (used as reference for background fluorescence) and P*_spoIIAA_ gfp* strains was monitored over time. 100 μL of the suspensions obtained in shaking flasks (as previously described) were distributed in microplates and measurements were performed with a microplate photometer (VICTOR™ X, PerkinElmer) equipped with an excitation filter at 485 nm and emission filter at 535 nm for green fluorescence measurement. The duration of the excitation was 1.0 s.

### The growth-sporulation model

The model of growth and sporulation can be divided into two modules. The vegetative cells’ growth was described by a classical primary model that has been previously developed: the modified logistic model of Kono (40) (Equation 1) and the sporulation kinetics were described from growth kinetics (Equation 2).

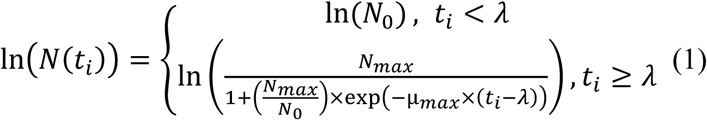

with *N_0_* the concentration of the inoculum (CFU/mL), *λ* the lag before growth (h), *μ_max_* the maximum vegetative growth rate (h^-1^), and *N_max_* the maximal concentration of total cells (CFU/mL). *N_max_* corresponds to the maximal concentration of vegetative cells reached at the stationary phase. Once the first spores appear, *N_max_* corresponds to the total cells, *i.e.* the spores and the remaining vegetative cells that have not differentiated into spores.

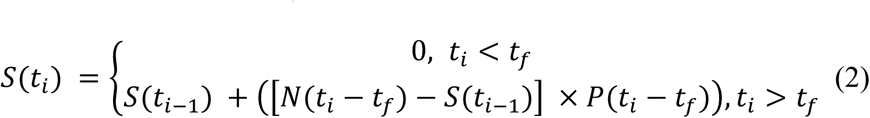

where *N*(*t_t_*— *t_f_*) are the total cells at time *t_i_* — *t_f_* given by equation 1, *S*(*t*_*i*-1_) are the spores at time *t*_*i*−1_ and *P*(*t_i_* — *t_f_*) is the probability of the vegetative cells committing to sporulation at time *t_i_* — *t_f_*.

The probability to commit to sporulation was defined as the proportion of cells that commit to sporulation over time. Previous works have shown that vegetative cells of a bacterial population do not initiate the sporulation at the same time (30). Consequently, the probability to sporulate evolves over time. In order to describe this evolution, four density functions (the Gaussian, the Weibull, the Lognormal and the Gamma laws) were evaluated and compared on four criteria: the biological significance of each-model parameters, the parsimonious number of parameters and the quality of fit of the kinetics with the RMSE statistical criterion (see below, equation 8). This led us to choose the Gaussian (or normal) probability density which was weighted by the maximal proportion *P_max_* of the vegetative cells to sporulate (equation 3).

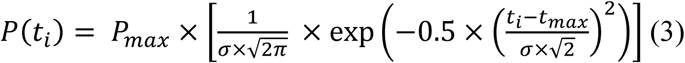

with *P*(*t_i_*) the probability of forming a spore at time *t_i_* (h^-1^), *P_max_* is the maximal proportion of vegetative cells forming spores (unitless). *P_max_* was obtained at the time *t_max_* (h) at which the cell has the maximal probability of initiating sporulation and *σ* the standard deviation around *t_max_* (h). Let us note that the maximal probability to sporulate at time *t_max_P(t_max_)* can be calculated as follows: 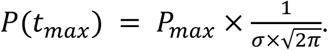 Finally, the sporulation module of the global model of growth and sporulation combines the equations 2 and 3.

### Methodology to assess the growth and the sporulation parameters

The growth and the sporulation parameters of the model in equations 1 and 4 were estimated in a three-step procedure.

In the first step, the primary growth model was fitted to the experimental counts (ln (CFU/mL)) to estimate the growth parameters (*N_0_ λ, μ_max_* and *N_max_*) with Equation 1.

In the second step, the experimental fluorescence data in log_10_ (AU) were plotted against time in order to estimate the mean time taken to initiate the sporulation (*t_max_*) and the probability scattering *σ*. We considered that within the population, each cell of strain P*_spoIIAA_ gfp* that commits to sporulation produces the same amount of GFP, *i.e.* has the same fluorescence intensity. A sporulating cell is composed of a mother cell and a forespore. The mature spore is released into the medium after lysis of the mother cell. Consequently, the fluorescence measured in a bacterial population corresponds to the fluorescence emitted by sporulating cells in addition to the fluorescence of the medium linked to the GFP molecules released in the medium following the lysis of the mother cell. To simplify the equations, the fluorescence that would be related to the presence of GFP molecules in the refractive spores is neglected. Consequently, the accumulation of fluorescence was directly related to the accumulation of cells that have initiated the sporulation and ultimately, to the accumulation of spores *i.e.* the sporulation kinetics.

The auto-fluorescence of the wild-type strain BSB1 was used as the background fluorescence. The two BSB1 and P*_spoIIAA_ gfp* strains were concomitantly cultivated. The fluorescence emitted by strain BSB1 was subtracted from the fluorescence emitted by strain P*_spoIIAA_ gfp* at each time point to assess the fluorescence associated with the production of GFP, hereafter referred as the “fluorescence”. The fluorescence kinetics were fitted with the cumulative distribution function for the normal distribution (equation 4). This function is used to assess the probability of a cell initiating the sporulation over time (equation 2 and 3 and Figure 1).

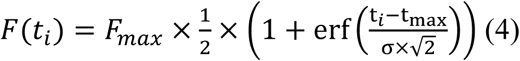

with *F*(*t_i_*) the fluorescence at time *t_i_* (AU), *F_max_* the maximal fluorescence (AU), *t_max_* (h) the time at which *F_max_* (UA) is obtained, *σ* the standard deviation around *t_max_* and erf, the error function of Gauss.

In the third step, the time taken to form a spore (*t_f_*) and the maximal proportion of sporulating *P_max_* were estimated: the sporulation curves were fitted with the Gaussian distribution function (equation 5) modified as follows:

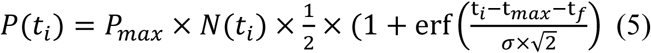

with *N*(*t_i_*) the concentration of total cells (equation 1), *t_max_* (h) the time at which *F_max_* (UA) was obtained, *P_max_* was the maximal proportion of sporulating cells, and *σ* (h) the standard deviation around *t_max_* (h). *P_max_* and *t_max_* were estimated in the previous step, by fitting the fluorescence kinetics in equation 5, and were used as inputs in equation 6 to fit the sporulation kinetics. The two parameters fitted on the sporulation kinetics were *P*_max_, and the time to form a spore *t_f_*.

### Statistical procedures and analysis

The growth and sporulation parameters of equations 1 to 6 were estimated by minimizing the Error Sum of Squares (ESS, fmincon, Optimization Toolbox; MATLAB 7.9.0; The Math-works, Natick, USA) (equation 6). 95% confidence intervals were estimated with the nlparci function of the Optimization Toolbox (MATLAB 7.9.0; The Math-works, Natick, USA).

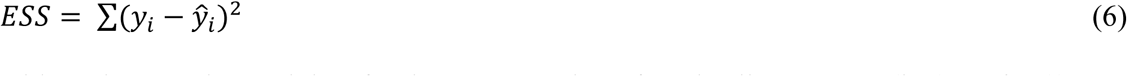

with *j_i_* the experimental data for the concentration of total cells or spores (ln (CFU/mL)) or fluorescence (AU) and *ŷ_i_* the value calculated with the model.

The goodness of fit of the model was assessed with the RMSE (Root Mean Square Error):

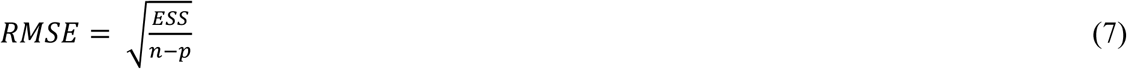

with ESS, the Error sum of squares calculated in equation 6, n, the number of experimental data and *p* the number of parameters of the model.

The likelihood ratio test (50) was used to check that the growth and sporulation kinetics were not significantly different between the wild type BSB1 and P*_spoIIAA_ gfp* strains. The growth and sporulation parameters were estimated for both strains. In order to compare the quality of fit with the model with fitted parameters or inputs, the likelihood ratio (S_L_) was calculated as follows (50):

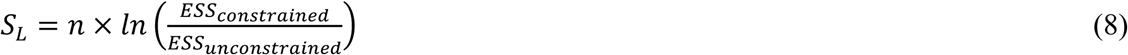

where *n* is the number of experimental data, *ESS_unconstrained_* is the ESS obtained by fitting the eight growth and sporulation parameters to the kinetics of the strain P *_spoIIAA_ gfp* and *ESS _constrained_* is the ESS obtained with the same eight kinetics but using the 8 parameters estimated on strain BSB1 as inputs. The value was compared with the Chi-squared value (15.51) that corresponds to a degree of freedom of eight and a tolerance threshold *α* of 5%.

## Acknowledgement

We thank Dr. Arnaud Chastanet (Micalis, Jouy-en-Josas, France) for providing the *B. subtilis* AC699 strain. This work was supported by Quimper Communauté and by a doctoral grant from Région Bretagne (France).

